# Structural Dynamics of LDL Receptor Interactions with E498A and R499G Variants of PCSK9

**DOI:** 10.1101/2024.07.25.605225

**Authors:** Nur Alya Amirah Azhar, Hapizah Nawawi, Yung-An Chua, Siti Azma Jusoh

## Abstract

The low-density lipoprotein receptor (LDLR) plays an integral role in cellular cholesterol uptake and lipid metabolism by primarily regulating hepatic clearance of plasma low-density lipoprotein cholesterol (LDL-C). Physiologically, proprotein convertase subtilisin/kexin type-9 (PCSK9) attenuates LDLR function by binding to the LDLR extracellular domain, leading to its lysosomal degradation and thereby preventing the total depletion of circulating LDL-C. However, pathogenic variants of PCSK9 are able to reduce the availability of LDLR, thus significantly increasing plasma LDL-C levels. Despite this understanding, the detailed molecular mechanism of LDLR-PCSK9 interaction remains elusive due to the lack of a full atomistic structure of LDLR. In this study, molecular dynamics (MD) simulations were employed to predict LDLR structural dynamics upon binding to PCSK9. Furthermore, two PCSK9 variants, E498A and R499G that were identified in clinically diagnosed Malaysian FH patients were investigated for their mutational effects. The simulations, spanning 500 ns, were conducted for three LDLR-PCSK9 complexes: LDLR-PCSK9 wild-type (WT), LDLR-PCSK9 (E498A), and LDLR-PCSK9 (R499G). Throughout the simulations, PCSK9 structure remained highly stable, in contrast to the LDLR structure that sampled large conformational space. The WT complex exhibited the least change, whereas the R499G complex displayed the most pronounced conformational rearrangement. During the simulations of WT and E498A complexes, the β-propeller domain of LDLR formed interactions with the prodomain of PCSK9. Aligned with the observation, the MM/GBSA analysis revealed that the E498A complex exhibited the highest LDLR-PCSK9 binding affinity (−63.81 kcal/mol), followed by the WT complex (−33.07 kcal/mol), and the R499G complex (−24.21 kcal/mol). These findings provide novel insights into the dynamic interactions between LDLR and PCSK9, highlighting the importance of structural flexibility in mediating their functional relationship. Further studies with complete LDLR structures are required to fully elucidate the molecular mechanisms underlying LDLR-PCSK9-mediated cholesterol homeostasis.

## 1. Introduction

The low-density lipoprotein receptor (LDLR), primarily present on the cell surface of liver cells, is crucial for the reuptake of serum low-density lipoprotein cholesterol (LDL-C). Synthesized in the endoplasmic reticulum, LDLR encoded by the LDLR gene is composed of 839 amino acid residues after the removal of a 21-amino acid signal peptide (Surdo et al., 2011). In humans, LDLR expression and LDL-C endocytosis are essential for maintaining cholesterol homeostasis and lipid metabolism, as LDL-C accounts for 75% of circulatory lipoprotein, with the majority of them being cleared by LDLR (Gu × Zhang, 2015). When LDLR quantity on the cell surface is low, LDL-C clearance is reduced, leading to elevated plasma LDL-C concentrations. Conversely, an abundance of LDLR increases LDL-C removal, resulting in decreased plasma LDL-C levels (Feingold, 2022; Pirahanchi et al., 2023). Pathogenic mutations in the *LDLR* gene can impair the protein’s function, leading to familial hypercholesterolemia (FH). Thus, normal LDLR structure and expression are vital for maintaining appropriate cholesterol levels and preventing excess LDL-C accumulation in the bloodstream (Ference et al., 2017). Untreated FH significantly increases the risk of premature cardiovascular disease (CVD). *LDLR* mutations are the primary cause of FH, accounting for 90-95% of FH-related mutations (Sun et al., 2018). The Leiden Open Variation Database (LOVD) database shows that the *LDLR* mutations have been comprehensively investigated, with over 4000 variants discovered (Leigh et al., 2017).

The mechanism of LDLR was first demonstrated by Brown and Goldstein in the 1970s through their research on cholesterol metabolism in *in vitro* fibroblasts isolated from FH patients (Goldstein & Brown, 1974). LDLR functions by recognizing apolipoprotein B (APOB) of LDL-C at neutral pH, triggering the internalization of the LDLR/LDL-C complex via receptor-mediated endocytosis into hepatocytes (Go × Mani, 2012; Surdo et al., 2011). In endosomes at acidic pH, LDLR releases LDL particles, recycles back to the cell surface, and binds to other LDL particles. Meanwhile, LDL particles are transported to lysosomes for degradation, and cholesterol is released (Pirahanchi et al., 2023). LDLR consists of an extracellular domain, a transmembrane domain, and a cytoplasmic domain (Rudenko et al., 2002; Yamamoto et al.,1984). The extracellular domain consists of a ligand-binding domain (LBD, with repeats L1-L7) and epidermal growth factor precursor homology domains (EGF(A) and EGF(B), a β-propeller, and an EGF(C) region) (Surdo et al., 2011). The LBD consists of ∼40 amino acid repeats, also known as LDL-receptor modules (Defesche, 2004). The second extracellular domain includes ∼400 amino acids, with two 40 amino acid repeats known as EGF(A) and EGF(B), a six-bladed β-propeller domain spanning 280 amino acids, and another EGF-like repeat, EGF(C) (Yamamoto et al.,1984). The EGF homology domain is crucial for LDL binding, LDL release at low pH in lysosomes, and LDLR recycling to the cell surface. The final ectodomain, called the O-linked polysaccharides domain, is located just outside the plasma membrane and consists of 58 amino acids (Rudenko et al., 2002; Surdo et al., 2011).

Proprotein convertase subtilisin/kexin type 9 (PCSK9) is a circulatory enzyme, primarily expressed in the liver, and plays a crucial role in regulating LDLR and consequently plasma LDL-C levels. PCSK9, encoded by the *PCSK9* gene translated to 692 amino acids that consist of a signal peptide, an N-terminal prodomain, a catalytic domain, and a C-terminal domain (Horton et al., 2009; Piper et al., 2007). Physiologically, PCSK9 prevents LDL-C clearance by binding and promoting LDLR degradation. The PCSK9-LDLR complex is internalized by endocytosis, and LDLR is degraded in lysosomes, decreasing LDLR levels on the cell surface (Horton et al., 2009). Therefore, PCSK9 has an LDLR-lowering effect, increasing plasma LDL-C levels (Xia et al., 2021). Many studies have identified pathogenic *PCSK9* variants as FH-causing genes, following *LDLR* and *APOB* (Abifadel et al., 2003; Brown & Goldstein, 1979; Innerarity et al., 1987). Gain-of-function (GOF) mutations in *PCSK9* lead to increased PCSK9 activity, resulting in excessive LDLR degradation (Hopkins et al., 2015; Sarkar et al., 2022). This reduction in LDLR on the cell surface causes elevated LDL-C levels, a hallmark of FH, which eventually leads to CVD.

The discovery of the LDLR-PCSK9 interaction identified the binding site between the catalytic domain of PCSK9 and the EGF(A) domain of LDLR (Zhang et al., 2007). Numerous studies reported on the interactions between the EGF(A) domain of LDLR and the catalytic domain of PCSK9 (Gu et al., 2013; Valenti et al., 2020), particularly examining the effects of point mutations at the binding interface. For instance, a study by Kwon et al., (2008) reported that the PCSK9 variant D374Y, located in the catalytic domain, increased the affinity of PCSK9 for LDLR, leading to decreased LDLR function and increased LDL-C levels in the blood. Additionally, researchers have developed antibodies targeting the catalytic domain of PCSK9 to block its binding to LDLR (Chan et al., 2009; Duff et al., 2009; Ni et al., 2011; Orringer et al., 2017), restoring LDL-C uptake. Therefore, the interaction between PCSK9 and LDLR is crucial as it influences LDLR function, underscoring its significance in lipid metabolism.

In this work, we employed molecular dynamics (MD) simulations to examine the dynamics of LDLR structure while interacting with PCSK9. Additionally, we investigated the impact of two novel PCSK9 variants, E498A and R499G, found in Malaysia FH patients (Razman et al., 2022) on the LDLR-PCSK9 complex and their interactions. The simulations captured the behavior and conformational changes of the LDLR structure while interacting with PCSK9.

## 2. Materials and Methods

### 2.1 Preparation of the LDLR-PCSK9 Complex Structures

The crystal structure of the LDLR-PCSK9 (PDB ID 3P5C) complex was retrieved from the Protein Data Bank (PDB) (Surdo et al., 2011). For the preparation as the starting input material, co-crystallized calciums were removed from the PDB file. The missing regions of the PCSK9 and LDLR complex were reconstructed using the SPDBV program (Guex & Peitsch, 1997). The protonation states of the LDLR-PCSK9 were determined at neutral pH using PROPKA 2.0 (Søndergaard et al., 2011).

### 2.2 Modeling of the PCSK9 Variants

The PCSK9 variants, E498A and R499G were discovered by the study of Razman et al., (2022). Both novel variants were identified among 35 clinically diagnosed Malaysian FH patients and classified as likely pathogenic according to ACMG guidelines (Richards et al., 2015). The model of the PCSK9 variants was developed using the SPDBV program. Including the wild-type complex, models of three LDLR-PCSK9 structure complexes were prepared as the input for the MD simulations (Figure 1).

**Figure 1.**
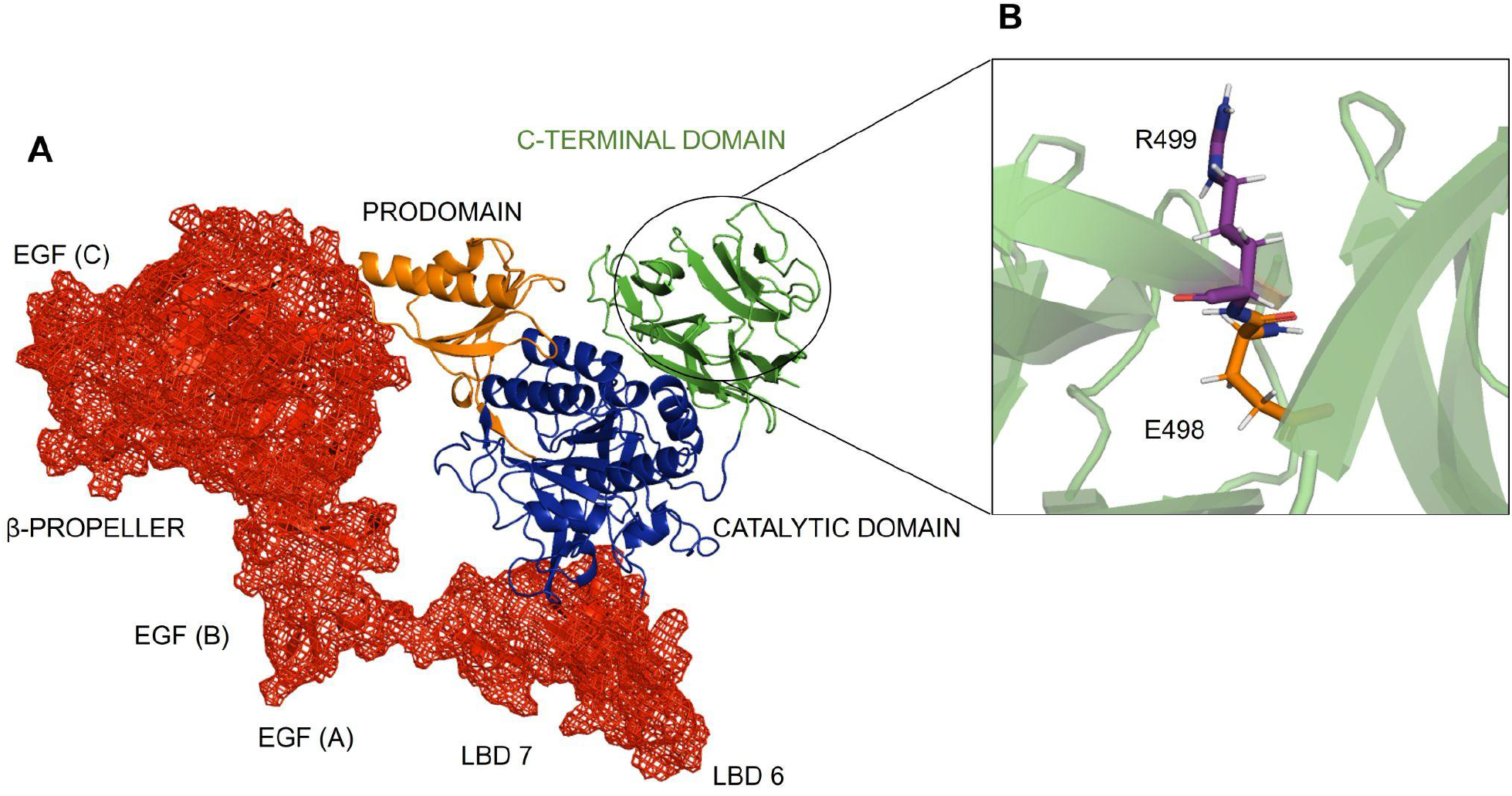
The X-ray crystallographic structure of the LDLR-PCSK9 complex (PDB ID: 3P5C). The structure represents PCSK9 and LDLR interactions, which was utilized as the initial structure for the modeling of *PCSK9* mutants (A). Two amino acid residues positioned at the C-terminal domain of PCSK9 (R499 and E498) were substituted with glycine and alanine, respectively, in order to model the structure of PCSK9 novel variants, which were discovered in clinically diagnosed Malaysian FH patients (Razman et al., 2022) (B).

### 2.3 Molecular Dynamics Simulations

For each PCSK9 variants-LDLR complex, a water box using TIP3P water molecules with edges of 12 Å from the protein in the simulation cubic box was added. The complex was later neutralized with NaCI concentration and solvated with ∼120,000 water molecules. Each system is composed of ∼ 350,000 atoms. The system setup for the wild-type LDLR-PCSK9 complex was prepared using the same procedure.

MD simulations were carried out using GROMACS 2023 (Abraham et al., 2015) with the CHARMM36M force field (J. Huang et al., 2017). Prior to the production simulations, the initial system was subjected to 1000 steps of energy minimization using the steepest descent algorithm to remove interatomic clashes. Following the minimization, an NVT equilibration (constant temperature) and NPT (constant pressure) equilibration were run. Hydrogen atoms were constrained using the LINC algorithm (Hess et al., 1997). The temperature of each complex was stabilized at 310.15 K for the protein and water/ions using a Nose-Hoover thermostat. Coulomb-long range interactions were evaluated using the Particle Mesh Ewald (PME) method with a cutoff set at 1.2 nm. For the van der Waals interactions, a cut-off distance of 1.2 nm was used. The simulations were carried out for 500 ns for each system with 2 fs timesteps under the isothermal-isobaric (NPT) ensemble, which maintains a constant number of particles, temperature, and pressure. For each system, simulations were performed in replications to ensure correct data productions.

### 2.4 Simulation Trajectory Analysis

Analyses of the root mean square deviation (RMSD), root mean square fluctuation (RMSF), and radius of gyration were carried out by GROMACS tools. The structural parameters which are RMSD and RMSF were measured using gmx_rms and gmx_rmsf, respectively. Hydrogen bonds were analyzed using the gmx_hbond tool. The output data were analyzed using the XMGrace program (https://plasma-gate.weizmann.ac.il/Grace/), and the plots of the analysis were prepared using Gnuplot (http://www.gnuplot.info/). The PyMOL (Schrödinger and DeLano, 2020) and VMD program (Humphrey et al., 1996) was used for visualization, hydrogen-bond interaction analysis, and image-making.

### 2.5 MM/GBSA calculation

Molecular Mechanics/Generalized Born Surface Area (MM/GBSA) was used to calculate the free binding energy between PCSK9 and LDLR of each complex based on the final 100 ns trajectories (400-500 ns) using gmx_MMPBSA (Valdés-Tresanco et al., 2021). Briefly, the free-binding energy can be expressed as:

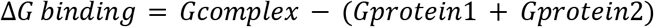

Where *Gcomplex* refers to the LDLR-PCSK9 complex, and protein 1 and protein 2 are the respective proteins in the complex. The individual free energies for each component above are determined by:

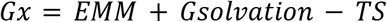

Where *EMM* is the molecular mechanics energy, *Gsolvation*, the solvation energy, and TS is the entropic contribution. The molecular mechanics energy and solvation energy can be further broken down into their component energies:

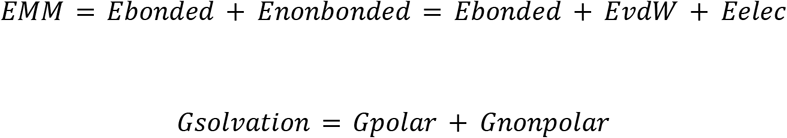

Here, *Ebonded* is zero, since we have used the single trajectory approach. *EvdW* and *Eelec* are the van der Waals and electrostatic contributions to the vacuum binding, respectively, while *Gpolar* and *Gnonpolar* are the electrostatic and non-electrostatic contributions to the solvation energy.

## 3. Results

### 3.1 Overall Structural Dynamics of LDLR-PCSK9 Complexes

MD simulations offer a valuable technique for investigating structural dynamics of proteins which allow us to gain insights into time-dependent changes. In this study, we employed MD simulations to understand the structural dynamics of LDLR in-complexed with PCSK9. In addition, we investigated the impact of two PCSK9 variants, E498A and R499G to the protein structure as well as the interactions between LDLR and PCSK9. Hence, we built three model systems of LDLR-PCSK9 complexes representing the LDLR-PCSK9 wildtype (WT complex), LDLR-PCSK9(E498A) (E498A complex), and LDLR-PCSK9(R499G) (R499G complex). Unbiased atomistic MD simulations were performed for a duration of 500 ns for each LDLR-PCSK9 complex.

The crystal structure of LDLR-PCSK9 used as a starting structure in this study exhibited that both proteins have interactions only between the EGF (A) region of LDLR and the catalytic domain of PCSK9. During the simulations, we observed in all systems that the overall structure of PCSK9 had no significant changes. Conversely, the LDLR structure underwent large conformational changes, specifically at the EGF(A) and EGF(B) regions causing the displacement of the β-propeller region. Despite the conformational changes, the EGF(A) region remained interacting with the catalytic domain of PCSK9. In addition, the incomplete LBD region of LDLR (LBD 6 and 7) was observed to have different behaviors in the wild type and the mutant systems (Figure 2). In the WT complex, the β-propeller region of LDLR was observed to immediately move away from the PCSK9 prodomain starting from the beginning of the simulations. Whereas, a residue segment of 623-626 shifted closer towards the prodomain, establishing interactions with a loop region (residue 73-76). During the ∼100 ns, another region of the β-propeller (residue 580-585) came in contact with the PCSK9 prodomain region (residue 105-110). (Figure 2A). Interactions between the β-propeller region of LDLR with the PCSK9 prodomain retained till the end of the 500 ns simulations, with various residues within the domains participating in the interactions.

**Figure 2.**
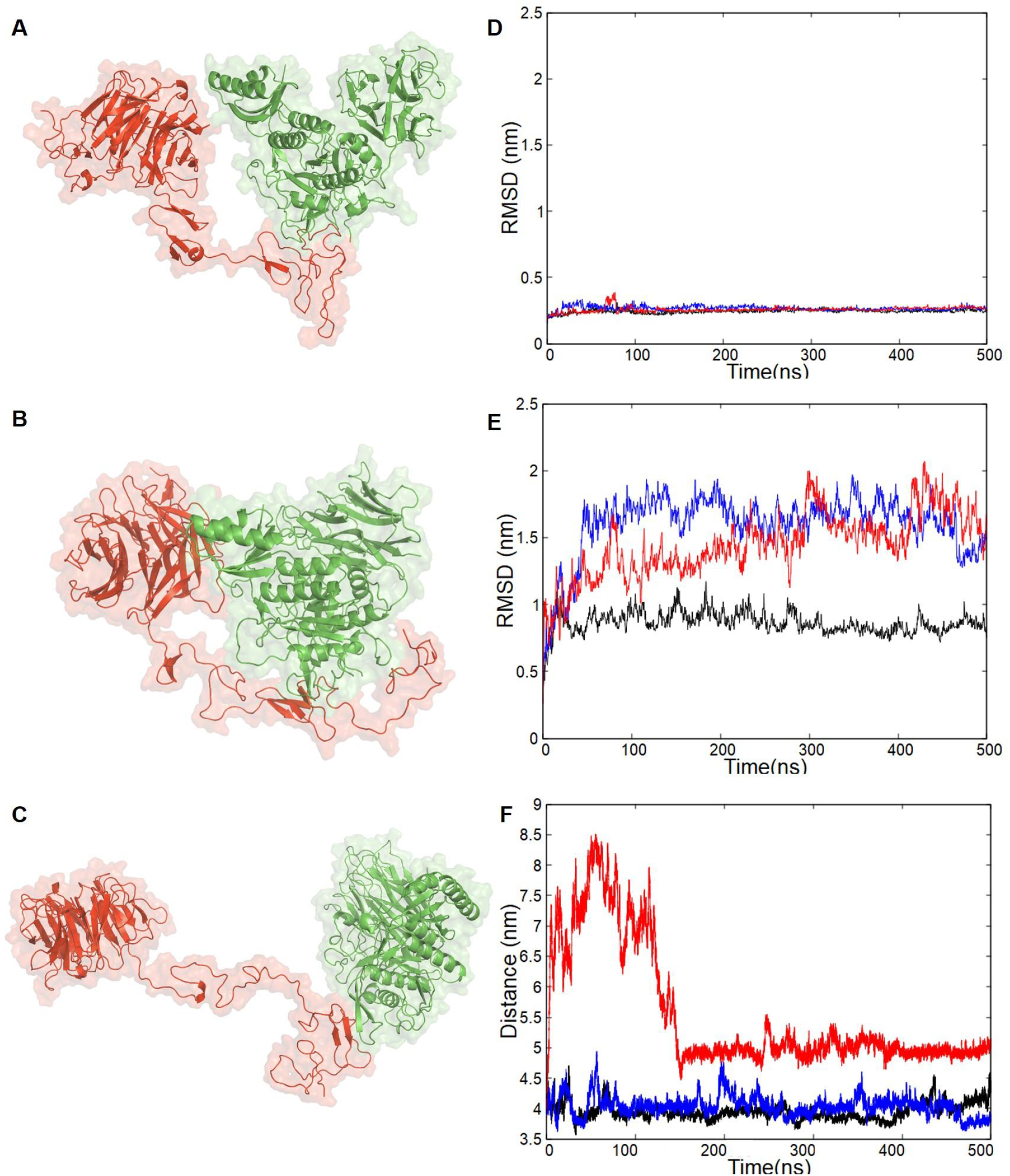
3D structure visualization and MD simulation analysis of LDLR-PCSK9 complex. Snapshots of WT complex (A), E498A complex (B), and R499G complex (C). RMSD of the PCSK9 (D) and the LDLR (E) during the 500 ns of MD simulations. Distance between the β-propeller of LDLR and prodomain of PCSK9 (F). Black (WT complex), blue (E498A complex), and red (R499G complex).

A similar trend was observed in the E498A complex, where regions of the β-propeller were interacting with the prodomain of PCSK9 throughout the simulations. A residue segment of 437-443 of the β-propeller formed interactions with residues 70-75 of the prodomain, facilitating proximity between LDLR and PCSK9 in these regions. Notably, in contrast to the WT complex, the initial structure of the E498A complex displayed no interactions with PCSK9. However, during the simulations, the regions comprising LBD 6 and 7 (residues 260-296) underwent from folded to unfolded state before 100 ns, and then later formed sustained interactions with the catalytic domain of PCSK9 (Figure 2B).

In the simulations of the R499G complex, the LDLR shows a distinct behavior of the β-propeller compared to the other two systems (Figure 2C). At the initial stage of the simulations (0-190 ns), the β-propeller region moved away from the prodomain of PCSK9 up to 8.5 nm distance (Figure 2F). From ∼200 ns onwards, the β-propeller maintained a distance of 5 nm from the prodomain but still did not form any interaction with PCSK9. In addition, the LBD 6 and 7 regions exhibited folding behavior and curled backward towards the EGF (A), and formed interactions within the regions. The lack of interaction between the β-propeller of LDLR and the prodomain caused the β-propeller to remain at a distance from the PCSK9 until the end of the simulations.

### 3.2 Root Mean Square Deviation (RMSD)

The dynamic structures of the LDLR-PCSK9 complexes were quantitatively analyzed using various metrics. One widely used measure, RMSD, assesses protein equilibration and structural stability by comparing each protein’s conformation to a reference structure. Differences from the reference indicate conformational changes. In this study, we employed RMSD analysis to evaluate the structural stability of both PCSK9 and LDLR in all three complexes (Figure 2D & 2E).

Throughout the simulations, PCSK9 exhibited minimal conformational changes when complexed with LDLR. The RMSD values for PCSK9 were 0.24 nm ± 0.01 for the WT complex, 0.27 nm ± 0.02 for the E498A complex, and 0.26 nm ± 0.20 for the R499G complex. These values indicate that the overall structure of PCSK9 remained largely unchanged during the simulation. In stark contrast, LDLR showed a greater degree of structural deviation. The RMSD for LDLR in the WT complex was 0.86 nm ± 0.08. Although there were slight fluctuations observed throughout the simulations, LDLR managed to retain its overall stability. However, the E498A complex showed a significantly higher RMSD of 1.60 nm ± 0.23, nearly double that of the WT. This suggests a considerable increase in structural deviation for LDLR when complexed with the E498A variant of PCSK9. Similarly, the R499G complex also showed an increased RMSD for LDLR, with a value of 1.45 nm ± 0.24. While this is slightly lower than the E498A complex, it is still notably higher than the WT. The elevated RMSD values for LDLR compared to PCSK9 highlight a substantial structural stability between the two protein molecules.

### 3.3 Radius of Gyration (Rg)

We conducted a Rg analysis for the LDLR-PCSK9 complexes. The Rg is a measure of the size of a protein molecule and can provide insights into the compactness of the protein structures. It is a particularly useful metric in molecular dynamics simulations as it can help assess the stability of protein folding. A protein that is stably folded will maintain a relatively consistent Rg over time, indicating that the protein remains compact and retains its structure. Conversely, a protein that is unfolding or fluctuating in its structure will exhibit changes in its Rg, reflecting the alterations in the protein’s shape and size.

The results show across all three complexes that PCSK9 displayed low and steady Rg values. This indicates stable structural dynamics and compactness within the simulations (Figure 3A). Specifically, the PCSK9 of the WT complex exhibited an Rg value of 2.59 nm ± 0.01. The PCSK9 structure in the E498A and R499G complexes showed similar stability, with Rg values of 2.60 nm ± 0.01 and 2.58 nm ± 0.02, respectively. In contrast, the LDLR in the complexes exhibited higher Rg values with higher standard deviations. This suggests conformational changes and structural flexibility within the complexes (Figure 3B). The LDLR of the WT complex had an Rg value of 3.31 nm ± 0.08. The E498A complex displayed an Rg of 3.50 nm ± 0.10, while the R499G complex exhibited an Rg of 3.69 nm ± 0.31. Notably, the R499G complex demonstrated the highest average Rg and the greatest deviations in Rg values. This suggests a higher degree of structural flexibility or instability in this complex. On the other hand, the E498A complex exhibited lower deviations compared to both the R499G and WT complexes, indicating a more stable structure.

**Figure 3.**
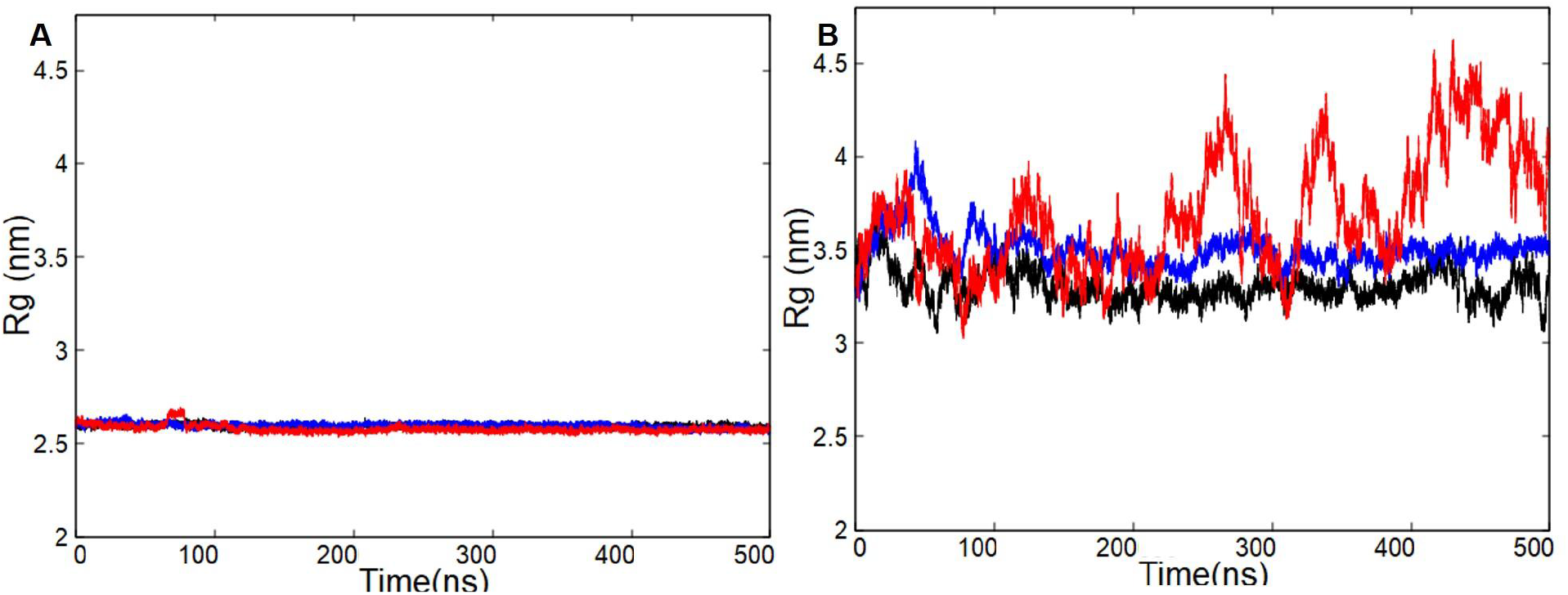
The Rg analysis of LDLR-PCSK9 complexes. The Rg values of PCSK9 (A) and LDLR (B) during the 500 ns of MD simulations. Black (WT complex), blue (E498A complex), and red (R499G complex).

### 3.4 Root Mean Square Fluctuation (RMSF)

RMSF analysis on the LDLR-PCSK9 complexes was aimed to identify residues that contribute significantly to the structural dynamics of both PCSK9 and LDLR. The analysis measures the average deviation of a particle over time from a reference position. It provides a quantifiable metric for the flexibility of individual residues in a protein structure during MD simulations. A residue maintaining a stable conformation will exhibit a relatively constant RMSF over time, indicative of a stably folded state. On the other hand, residues that are part of conformational changes or exhibit structural flexibility will have higher RMSF values. This is a reflection of a greater positional variability of the particular residue throughout the simulation.

In line with the previous analysis, the overall RMSF values for PCSK9 were significantly lower than those for LDLR, indicating that PCSK9 residues experienced minimal fluctuations across all three complexes (Figure 4A & 4B). This pattern suggests that PCSK9 maintained a stable conformation throughout the MD simulations. Based on the individual complexes, it was observed that PCSK9 residues in the the E498A complex showed only a small subset of residues exhibiting slight fluctuations, specifically at residue regions of 164-170, 211-222, 448-454, and 571-584. Meanwhile, in the R499G complex, the PCSK9 residues showed fluctuations across a broader range of residues, including 164-170, 192-227, 449-454, 490-518, 569-587, and 614-621. Despite the observed fluctuations in certain residues, PCSK9 structures demonstrated minimal conformational changes throughout the simulations. In stark contrast, the RMSF values of LDLR were significantly higher in all three complexes (Figure 4D). In each complex, the LDLR structure underwent high flexibility, especially in the R499G complex. Particularly notable were the fluctuations in residues 250-400, representing LBD 6 and 7, and the EGF(A) and EGF(B) domains. Meanwhile in the E498A complex, LDLR residues also displayed significant fluctuations in residues 250-375, covering LBD 6 and 7, EGF(A) and EGF(B), and residues 641-678 in the EGF(C) domain. These fluctuations indicate a higher propensity for conformational changes in LDLR compared to PCSK9. Overall, the RMSF analysis underscores the differential flexibility within the LDLR-PCSK9 complexes, highlighting the greater dynamic nature of LDLR as opposed to the more stable structure of PCSK9.

**Figure 4.**
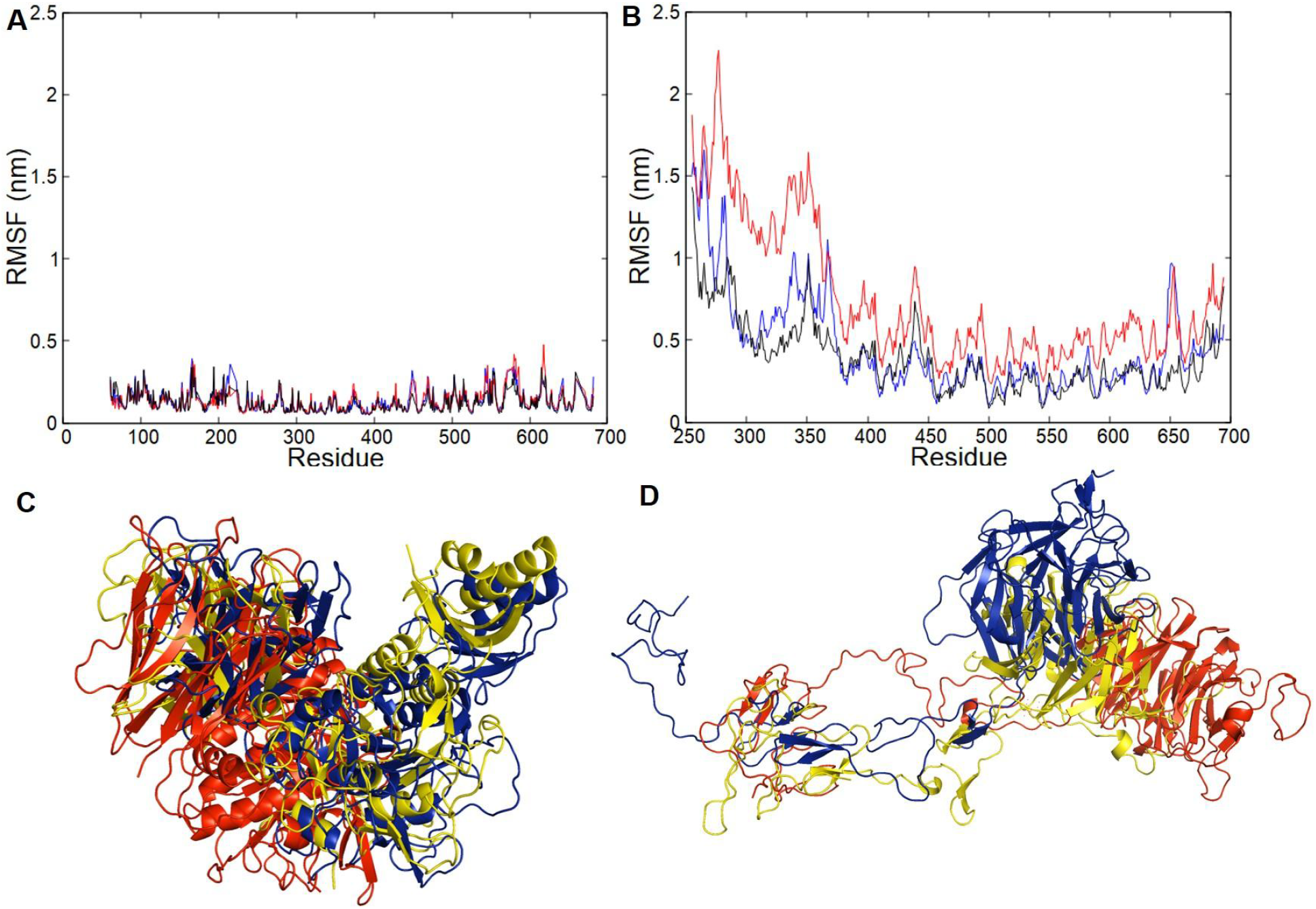
RMSF analysis and structural superimpositions of PCSK9 and LDLR. The RMSF analysis of PCSK9 (A) and LDLR (B) structures during the 500 ns of MD simulations. Black (WT complex), blue (E498A complex), and red (R499G complex). Superimposition of the PCSK9 (C) and LDLR (D) structures based on the 500 ns simulation snapshots. Cartoon representations of the protein structures are for the E498A complex (blue), R499G complex (red) and WT complex (yellow).

### 3.5 Molecular Mechanics Generalized Born Surface Area (MM/GBSA)

The MM/GBSA method is a computational approach employed to calculate the binding free energy of biomolecular complexes. This technique decomposes the binding energies into several components, including van der Waals interactions, electrostatic interactions, and solvation-free energies. This breakdown facilitates a deeper understanding of the stability and affinity of protein complexes. Here, we utilized the MM/GBSA method to assess the binding energies of the three LDLR-PCSK9 complexes (Table 1). We observed that the components contributing to the total binding affinity varied among each complex.

**Table 1.** Average binding energy calculation of PCSK9-LDLR complexes using MM/GBSA.

| LDLR-<br>PCSK9<br>complexes | Average binding energy (kcal/mol) |  |  |  |  |  |  |
| --- | --- | --- | --- | --- | --- | --- | --- |
| | $\Delta$ van der<br>Waals | $\Delta$<br>electrostatic | $\Delta$ GB | $\Delta$ Surface | $\Delta$ Gas | $\Delta$ Solvation | $\Delta$ Total |
| Wild-type | -75.69 | 323.57 | -270.03 | -10.92 | 247.88 | -280.95 | -33.07 |
| E498A | -104.00 | 124.22 | -68.93 | -15.09 | 20.21 | -84.02 | -63.81 |
| R499G | -42.18 | 523.64 | -499.51 | -6.16 | 481.45 | -505.67 | -24.21 |

MM-GBSA analysis of 500 ns MD simulations sheds light on the impact of mutations on PCSK9 binding to LDLR. The E498A mutation emerged as potentially favorable, evident from its most negative binding free energy (ΔG_bind, -68.9 kcal/mol) compared to both WT (−270 kcal/mol) and R499G (−499.5 kcal/mol). This advantage is further emphasized by the most negative van der Waals energy (−104 kcal/mol) in the E498A complex, indicating stronger attractive forces between the PCSK9 variant and LDLR. Interestingly, the E498A complex also had a lower electrostatic energy (124.2 kcal/mol) compared to the WT and R499G complexes, potentially reflecting reduced unfavorable interactions. However, a slightly less negative solvation energy (−84.02 kcal/mol) in the E498A complex suggests a potential trade-off, where the mutation might slightly affect solvation. Further supporting tighter binding, the E498A complex exhibited the most negative surface energy (−15.1 kcal/mol) compared to the WT (−10.9 kcal/mol) and R499G (−6.16 kcal/mol), indicating a larger buried surface area at the protein-protein interface, which is inline with the structural analysis. Conversely, the R499G complex stands out with the least favorable ΔG_bind, the least negative van der Waals energy, and a less negative surface energy, suggesting a significantly weaker interaction with LDLR, potentially due the large conformational changes in LDLR as observed structurally.

### 3.6 Structural Analysis of LDLR-PCSK9(E498A) and LDLR-PCSK9(R499G) Variants

In this study, we evaluated the impact of PCSK9 variants on their structure and interactions with LDLR. The selected variants are two novel PCSK9 variants identified in Malaysian familial hypercholesterolemia (FH) patients (Razman et al., 2022). The gene variants, PCSK9 c.1493A>C (E498A) and c.1495C>G (R499G), have evidence for pathogenicity according to the ACMG guidelines and are predicted to cause damaging effects by *in silico* prediction programs such as Polyphen and SIFT. These predictions support the functional relevance of the variants, as they reside in the C-terminal region known to interact with LDLR in endosomal low pH environments (Nozue, 2017). The study by Razman et al., 2022 virtually replaced the mutant residues to the X-ray structure of PCSK9 (PDB ID: 3P5C) to analyze potential disruptions to the interactions among the nearby residues. The prediction showed that the wild-type E498 formed hydrogen bonds with both S488 and S564. However, when the glutamic acid was mutated to the small, hydrophobic alanine (A498), it lost the interaction with S488. Similarly, the positively charged WT R499 participated in ionic interactions with a glutamic acid (E501) and also formed a hydrogen bond with an arginine (R510). When substituted with a glycine (G499), the mutant was predicted to lose the interaction with E501 and retained the hydrogen bonding with R510.

We employed MD simulations to investigate these interactions in a more dynamic setting, where the WT structure and both variant models of PCSK9 in complexes with LDLR underwent 500 ns duration of simulations. The results for the wild-type PCSK9 indicated the same interactions of E498 to S564 and S488, which were emphasized by a high number of hydrogen bonds (Figure 5A & 5C) as well as the interaction occupancy throughout the 500 ns of the simulations (55.87% and 21.75%, respectively). Meanwhile, the simulations also revealed other interactions, with the WT E498 also participating in hydrogen bonds with additional residues which are S563 and H597 (Supplementary Table 2). In line with these predictions, the MD simulations for the E498A variant also observed the predicted loss of interaction with S488. That could reflect the half reduction of hydrogen bonds compared to the WT. In addition, A498 was observed to interact with S564 and S563, which was not predicted from the previous findings (Figure 5B & 5D).

**Figure 5.**
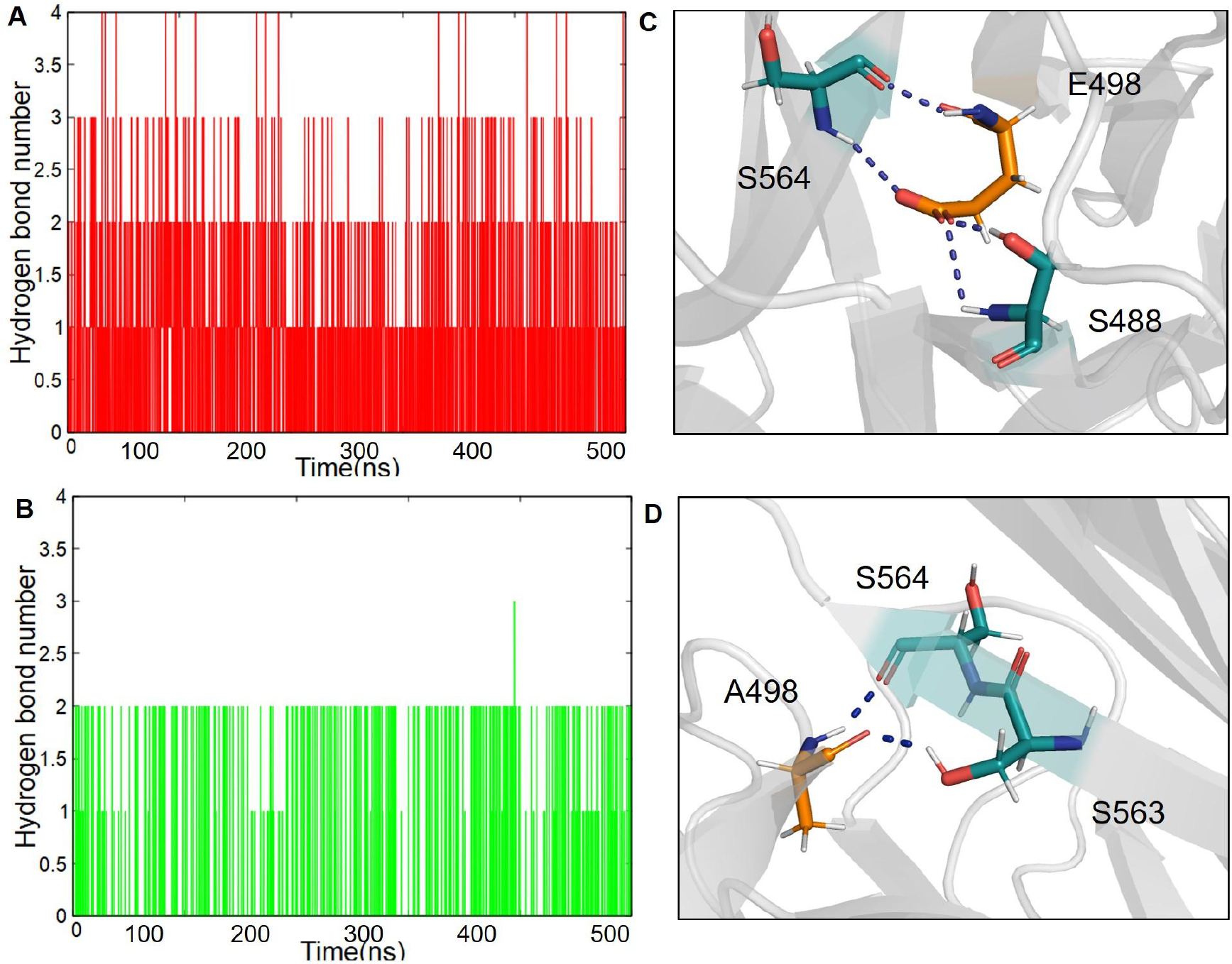
Hydrogen bond and interactions analyses of the WT and mutant residues of PCSK9(E498A). Number of hydrogen bonds of the WT E498 (A) and the mutant A498 (B) residues with their neighboring residues. 3D structure visualizations of the WT E498 (C) and the mutant A498 (D) interacting with the neighboring residues. PCSK9 structure (cartoon), WT and mutant residues (labeled stick), hydrogen bonds (dashed lines).

Similarly, the MD simulations indicated the R499G variant aligned with the prior predictions of a loss of interaction with E501. The WT R499 residue exhibited a broader repertoire of interacting with three residues (E501, R510, H565) compared to the G499 variant, which only formed polar interactions with R510 (Figure 6A & 6B). Interestingly, the simulations revealed a more extensive hydrogen bonding network for the wild-type R499 than previously anticipated that indicated R499 interactions with many neighboring residues. In the case for the mutant residue, G499, the simulations revealed its interactions with T513 (46.94%), R510 (44.57%), and D630 (33.44%) (Supplementary Table 3). This consistency highlights the potential of both MD simulations and structural analysis for predicting changes in hydrogen bonding upon mutation.

**Figure 6.**
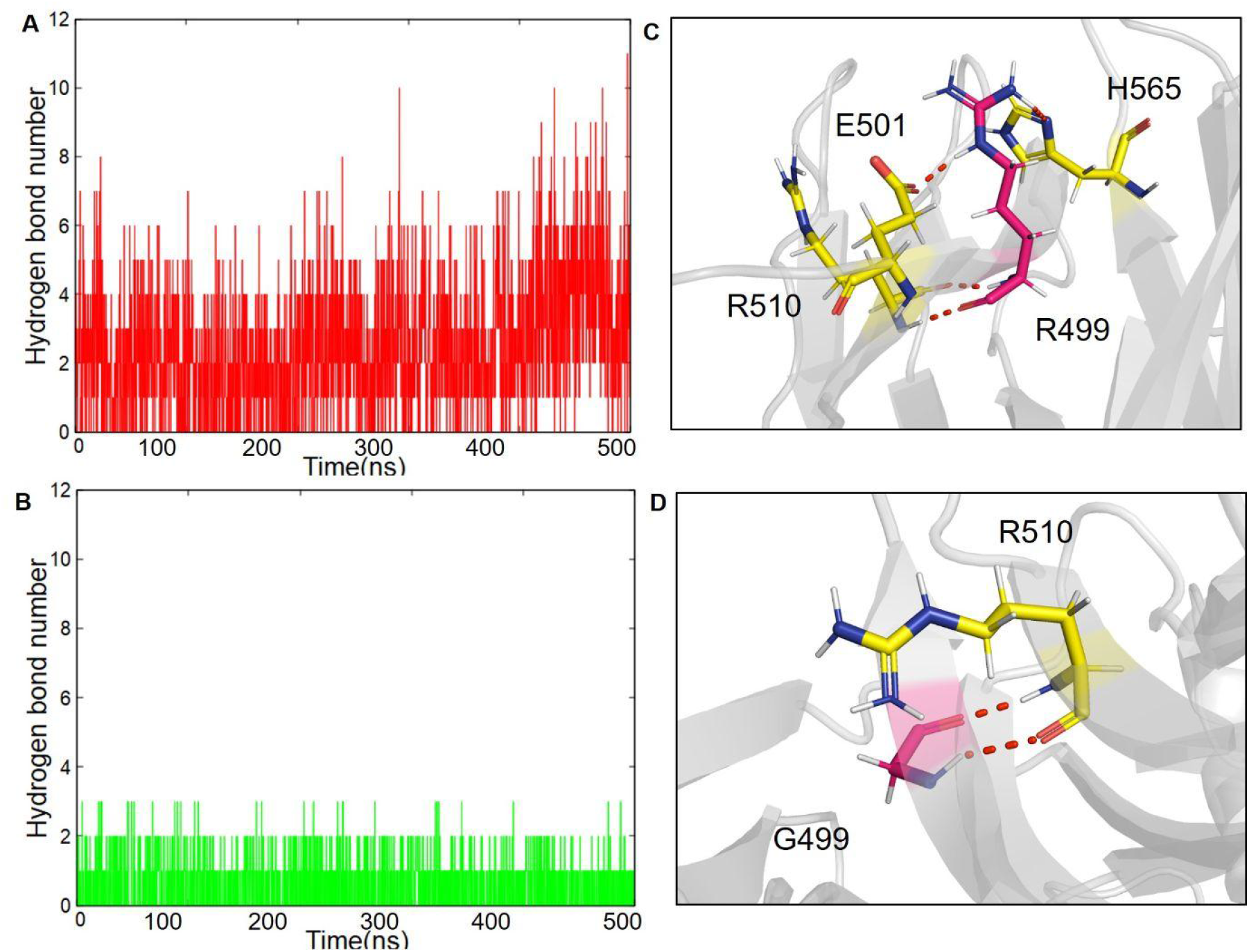
Hydrogen bond and interactions analyses of the WT and mutant residues of PCSK9(R499G). Number of hydrogen bonds of the WT R499 (A) and the mutant G499 (B) residues with their neighboring residues. 3D structure visualizations of the WT R499 (C) and the mutant G499 (D) interacting with the neighboring residues. PCSK9 structure (cartoon), WT and mutant residues (labeled stick), hydrogen bonds (dashed lines).

Overall, the MD simulations corroborated the key predictions regarding the impact of mutations on core interactions (loss of interaction with S488 for E498A and E501 for R499G). However, the MD simulations also revealed a more complex picture with additional hydrogen bonding partners for the wild-type residues, underlining the importance of considering dynamic behavior when studying protein-protein interactions.

## 4. Discussion

The primary function of LDLR is to efficiently remove LDL-C from circulation, preventing an increase in serum LDL-C levels (Benjannet et al., 2004; Kwon et al., 2008). On the cell surface, LDLR adopts an open conformation, crucial for its high-affinity binding to LDL-C (S. Huang et al., 2010). Upon endocytosis and transport to the endosome, LDLR undergoes a conformational change to a closed state due to the acidic environment. In this state, the LBD folds back onto the β-propeller, forming a loop-like structure with LBD 4 and 5 bound to the β-propeller (Rudenko et al., 2002). This conformational shift allows LDLR to release LDL-C particles for degradation before the receptor is recycled back to the cell membrane (Martínez-Oliván et al., 2015; Tveten et al., 2012). The structural flexibility of the LBD regions largely plays a major role in LDLR adopting an open conformation on the cell membrane and a closed conformation in hepatocytes. Contrary to LDL-C, PCSK9 does not bind to the LBD domain. The catalytic domain bind to LDLR EGF(A) domains, respectively in such a way to hold LDLR in an extended version, in order to prevent the LDLR rearrangement for recycling (Kwon et al., 2008; Surdo et al., 2011). However, a complete molecular understanding of the LDLR transition states as well as its interactions with PCSK9 remains unclear due to incomplete experimental structures of LDLR. The available experimental structures of LDLR lacked the sugar-rich domain, transmembrane domain (TMD), and cytoplasmic domain (Klee & Zimmermann, 2019; Rudenko et al., 2002). The longest X-ray structure of LDLR is in a closed conformation (PDB ID: 1N7D) (Rudenko et al., 2002) that consists of the seven LBD domains followed by EGF(A), EGF(B), B-propeller and EGF(C), yet still missing the other regions. Yet in this study, we utilized the shorter X-ray structure of LDLR, which represents the open conformation while interacting with PCSK9 (PDB ID: 3P5C) on the cell surface (Surdo et al., 2011).

MD simulation is a powerful tool in the field of structural biology, providing valuable insights into the dynamic nature of proteins. Unlike static experimental structures, MD simulations allow us to investigate the fluctuations and conformational changes of a protein. This dynamic information is crucial for understanding how protein flexibility contributes to their function. Moreover, MD simulation can provide deeper understanding of binding mechanisms, binding affinities, and the specific residues involved in complex formation (Badar et al., 2022; Hollingsworth & Dror, 2018). In this study, MD simulations were employed to investigate the dynamic behavior of PCSK9 and LDLR. Based on the 500 ns duration of simulations, the PCSK9 structure was highly stable, meanwhile LDLR exhibited significant conformational changes. However, the inherent flexibility of LDLR aligns well with the proposed mechanism of action of LDLR which undergoes a conformational shift upon LDL-C binding and subsequent release that required transition between open and closed states. On the other hand, the stability of PCSK9 suggests a more rigid structure, potentially crucial for its role in inhibiting LDLR function.

The simulations revealed significant structural rearrangements within the EGF and LBD domains of the LDLR, which may have caused the large movement of the β-propeller domain, yet its overall structure remained relatively stable. Notably in all complexes, the EGF(A) of LDLR maintained their interaction with the catalytic domain of PCSK9, consistent with established crystallographic structures and functional studies (Kwon et al., 2008; Surdo et al., 2011). However, in the E498A complex, LBD 6 and 7 were observed to interact with the catalytic domain of PCSK9. This condition is not reported elsewhere. Nevertheless, LBD 6-7 did not interact with the catalytic domain of PCSK9 in the WT and R499G complexes. The established reports based on *in vitro* studies demonstrated that the LBD of LDLR formed interactions to the PCSK9 C-terminal domain (Cunningham et al., 2007; Yamamoto et al., 2011). However, because the LDLR structure lacks LBD 1-5, the potential interaction between the LBD and the C-terminal cannot be observed, as LBD cannot reach the C-terminal. However, we may predict this possible behavior from the observed interactions between the LBD and the catalytic domain with future complete LDLR structure.

The X-ray structure of LDLR-PCSK9 structure exhibits that the β-propeller domain of LDLR forms van der Waals interactions with the prodomain of PCSK9 (Surdo et al., 2011). In this study, we observed stronger interactions between the β-propeller and the prodomain in the WT and E498A complexes based on hydrogen bond interactions. Conversely, in the R499G complex, the β-propeller domain, along with the EGF(B) and EGF(C) domains, moved away from PCSK9. The observed large conformational changes in the LDLR during the simulations might be attributed to the incomplete structure lacking the LBD regions, sugar-rich domain, and transmembrane domain (TMD). However, as the open state of LDLR facilitates LDL-C binding, the extended structure likely explores a large conformational space. The functional flexibility of LDLR and its interactions with other molecules are supported by previous reports. These include the interactions between the negatively charged LBD and positively charged APOB on LDL particles (Davis et al., 1987), the sugar-rich domain serving as a glycosylation site (Feingold, 2022), and the TMD anchoring the LDLR to cell membranes (Yamamoto et al., 1984).

This study also explored the dynamic interaction between LDLR and PCSK9 using MD simulations. Understanding these interactions is crucial in the context of FH, a genetic disorder characterized by elevated LDL-C levels. Genetic testing for FH variants plays a vital role in early diagnosis and risk assessment for CVD. Our simulations explored the structural effects of two novel PCSK9 variants (E498A and R499G) identified in clinically diagnosed Malaysian FH patients. These variants reside in the C-terminal region of PCSK9 and did not directly interact with LDLR throughout the simulations. However, the mutant complexes exhibited distinct conformational changes in the LDLR compared to the wild-type complex. These changes could be indirect effects arising from the PCSK9 variants, or they might be due to the limitations of the incomplete LDLR structure used in the simulations. Hence, it is important to consider established knowledge about PCSK9 variants.

Reports suggest that GOF mutations can increase PCSK9 activity towards LDLR, leading to increased LDLR degradation and decreased LDL-C uptake (Abifadel et al., 2003; Peterson et al., 2008). Conversely, loss-of-function (LOF) mutations decrease PCSK9 activity, resulting in less LDLR degradation (Berge et al., 2006; Cohen et al., 2006). Furthermore, studies have shown that a more positively charged C-terminal region of PCSK9 correlates with higher activity towards the negatively charged LBD region of LDLR (Deng et al., 2019; Tveten et al., 2012; Yamamoto et al., 2011). For instance, mutations replacing negatively charged glutamic acids with neutral glycine (E498A) or positively charged arginine with neutral alanine (R499G) could alter the overall net charge of the C-terminal domain. In theory, E498A might increase the positive charge, potentially enhancing binding between LDLR and PCSK9. Conversely, R499G could lead to a more negative charge, potentially weakening the interaction. However, the LDLR structure in this study lacked LBD regions 1 to 5. This limited structure might be too short to reach the C-terminal domain of PCSK9, explaining the absence of observed interactions between the variant residues (E498A and R499G) and the LBD of LDLR. Therefore, due to the absence of direct interactions between the variant residues (E498A and R499G) and the LBDs, there is no definitive relation of these mutations to the established GOF or LOF effects observed in experimental studies. Future simulations employing a complete LDLR structure are necessary to fully elucidate the functional consequences of these novel C-terminal PCSK9 variants in the context of FH.

## 5. Conclusion

This study investigated dynamics interactions of the LDLR-PCSK9 complex and structural impact of two genetic variants, PCSK9(E498A) and PCSK9(R499G). Our findings revealed a relatively stable PCSK9 structure in contrast to the highly dynamic nature of LDLR. The observed flexibility of LDLR suggests its conformational adaptability, which is likely crucial for its diverse functions, including the transition between open and closed conformations. MD simulations indicated disruptions in the residue-residue interaction network surrounding the mutated sites in the E498A and R499G variants, potentially influencing the PCSK9-LDLR binding interface. However, the absence of complete LDLR structural information limits our ability to fully assess the impact of these mutations. Understanding the intricate interplay between PCSK9 and LDLR is crucial for elucidating the mechanisms underlying familial hypercholesterolemia. This study provides insights into the potential effects of PCSK9 variants on LDLR-PCSK9 interactions. Further studies incorporating a complete LDLR structure and exploring the impact of different physiological conditions are necessary for a comprehensive understanding of this complex system.

## Supporting information

https://docs.google.com/document/d/1Q3OrgJGkPWoZxexHIhDBeG-eFWbzfYBFs3RB7BGuI_c/

## ACKNOWLEDGEMENTS

We acknowledge the Faculty of Pharmacy, UiTM Puncak Alam for the computing facilities used for this project.

## Author Contributions

NAAA: performed the experiments, analysis, and wrote the original draft. SAJ: Project administration, conceptualization, supervision, and approval of the final draft. YAC and HN: Manuscript review and editing. All authors have read and agreed to the published version of the manuscript.

## Funding

This study was funded by Universiti Teknologi MARA (UiTM) DUCS CoE Grant [600-UiTMSEL (PI. 5/4)(030/2022)].

## Conflicts of Interest

The authors declare no conflict of interest.

## Notes

### Competing Interest Statement

The authors have declared no competing interest.

