## Supplementary material for "Structural Dynamics of LDL Receptor Interactions with E498A and R499G Variants of PCSK9": https://docs.google.com/document/d/1Q3OrgJGkPWoZxexHIhDBeG-eFWbzfYBFs3RB7BGuI_c/

Supplementary materials

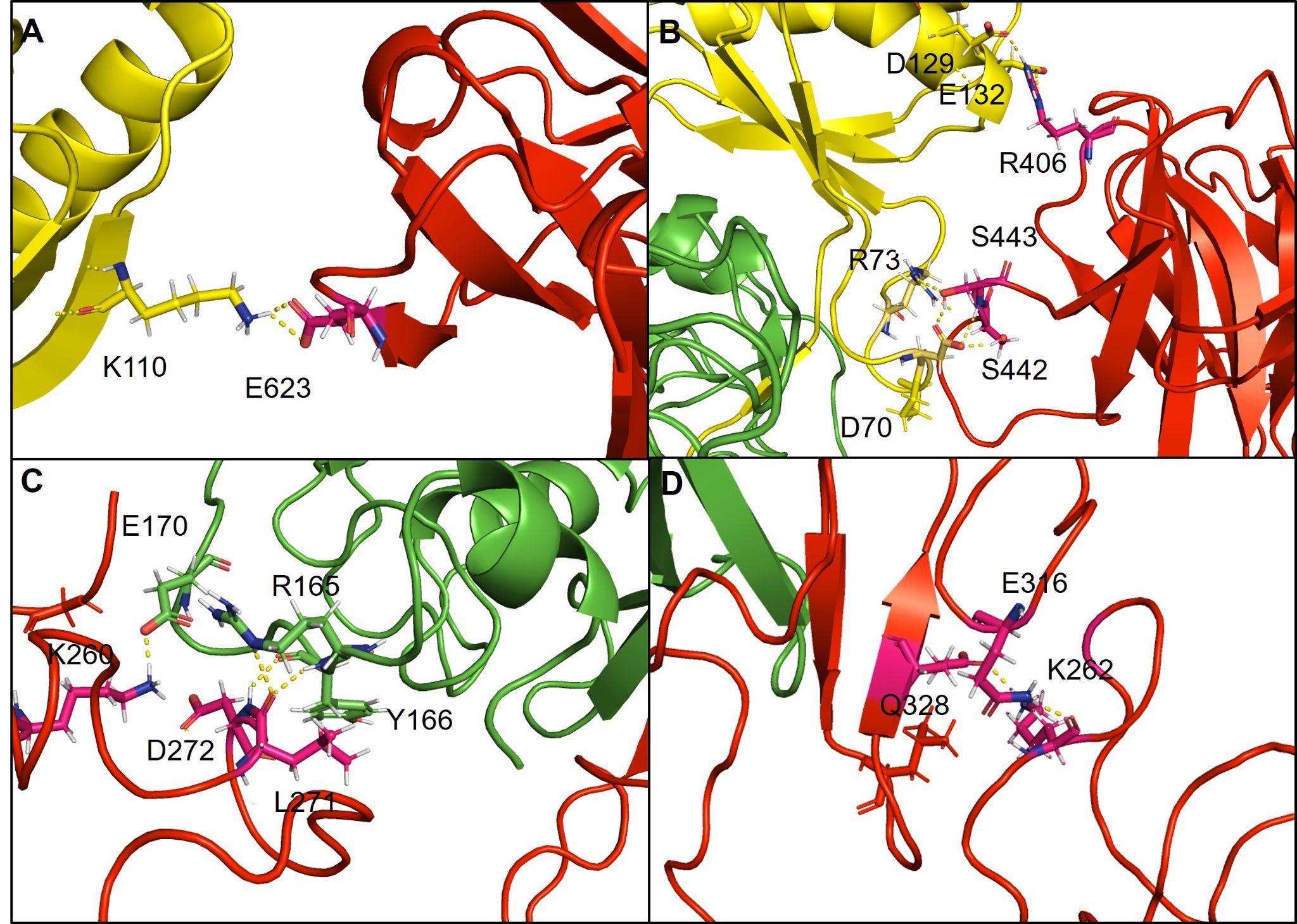

**Supplementary Figure 1 Snapshot of hydrogen bond interactions formed between LDLR and PCSK9 at 500 ns MD simulations.** The β-propeller residues interact with prodomain residues in the WT (A) and E498A complexes (B). The interaction formed between the LBD regions and catalytic domain in the E498A complex (C). The folding of the LBD region of the R499G complex causes the EGF (A) domain to form interactions within the region (D). Green cartoon (catalytic domain), yellow cartoon (prodomain), red cartoon (LDLR), WT and mutant residues (labeled stick) and hydrogen bonds (dashed lines).

**Supplementary Table 1** Hydrogen bond occupancy (%) between LDLR and PCSK9 residues within 3 Å of the LDLR-PCSK9 complex during 500 ns simulations.

| **WT complex** | | |
| --- | --- | --- |
| **PCSK9 residue** | **LDLR residue** | **H-bond occupancy (%)** |
| F379-Main | C308-Main | 60.11% |
| T377-Side | N309-Side | 56.07% |
| R194-Side | D310-Side | 21.79% |
| T377-Main | D310-Main | 45.98% |
| S153-Main | D299-Side | 52.3% |
| S381-Side | N301-Side | 2.92% |
| E332-Side | C255-Main | 10.33% |
| C331-Main | C255-Main | 1.40% |
| D333-Side | C255-Main | 3.28% |
| D238-Side | N295-Side | 1.36% |
| K125-Side | E623-Side | 21.23% |
| S127-Side | E605-Side | 2.88% |
| K110-Side | E623-Side | 30.68% |
| D367-Side | N301-Side | 2.80% |
| S381-Side | N301-Main | 28.15% |
| R104-Side | E581-Side | 7.41% |
| G106-Main | N604-Side | 8.73% |
| L108-Main | N604-Side | 18.94% |
| K125-Side | N624-Side | 13.02% |
| S153-Side | N295-Side | 1.56% |
| S153-Main | N295-Side | 1.88% |
| **E498A complex** | | |
| **PCSK9 residue** | **LDLR residue** | **H-bond occupancy (%)** |
| F379-Main | C308-Main | 62.52% |
| T377-Side | N309-Side | 57.51% |
| S381-Side | N301-Side | 4.77% |
| R194-Side | D310-Side | 4.77% |
| S153-Main | N295-Main | 3.12% |
| S153-Main | T294-Side | 1.52% |
| T377-Main | D310-Main | 35.20% |
| S153-Main | E296-Side | 4.97% |
| E267-Side | C255-Main | 3.04% |
| C268-Main | C255-Main | 7.01% |
| R215-Side | E351-Side | 2.52% |
| N295-Side | D238-Side | 1.36% |
| D70-Side | S443-Main | 12.90% |
| D70-Side | S442-Side | 46.42% |
| D70-Side | S443-Side | 17.74% |
| SER153-Side | D299-Side | 4.37% |
| D129-Side | R406-Side | 30.88% |
| E132-Side | R406-Side | 11.77% |
| D280-Side | C255-Main | 1.44% |
| S153-Main | D299-Side | 35.68% |
| D238-Side | K290-Side | 1.84% |
| R237-Side | E287-Side | 1.40% |
| R199-Side | E287-Side | 2.96% |
| E170-Side | K273-Side | 1.48% |
| S381-Side | N300-Side | 1.00% |
| R73-Side | V441-Main | 20.42% |
| D169-Side | K260-Side | 6.85% |
| E170-Side | K260-Side | 18.50% |
| A168-Main | D272-Side | 2.40% |
| Y166-Main | L271-Main | 30.44% |
| Y166-Main | D272-Main | 6.09% |
| R165-Side | E267-Side | 18.74% |
| R167-Side | E256-Side | 6.21% |
| S447-Side | E256-Side | 5.01% |
| D238-Side | T294-Side | 1.44% |
| R165-Side | E256-Side | 8.61% |
| D169-Side | K273-Side | 2.16% |
| D367-Side | N301-Side | 2.76% |
| S381-Side | N301-Main | 5.77% |
| D367-Side | S305-Side | 5.81% |
| S381-Main | H306-Main | 10.37% |
| R73-Main | V441-Main | 1.80% |
| R73-Side | S443-Side | 2.52% |
| D272-Side | C255-Main | 6.01% |
| R165-Side | T270-Main | 3.40% |
| ARG165-Side | L271-Main | 1.44% |
| E170-Side | G257-Main | 2.56% |
| **R499G complex** | | |
| **PCSK9 residue** | **LDLR residue** | **H-bond occupancy (%)** |
| F379-Main | C308-Main | 66.28% |
| R194-Side | D310-Side | 4.16% |
| S153-Main | D299-Side | 47.50% |
| T377-Main | D310-Main | 48.42% |
| S381-Side | N301-Side | 3.92% |
| T377-Side | N309-Side | 51.58% |
| D238-Side | T294-Side | 2.96% |
| S153-Side | D299-Side | 4.61% |
| R659-Side | E623-Side | 3.00% |
| D367-Main | N301-Side | 5.53% |
| S381-Main | H306-Main | 6.85% |
| R319-Side | E351-Side | 3.60% |
| A327-Main | C255-Main | 4.81% |
| D272-Side | C255-Main | 12.90% |
| S153-Main | N295-Side | 1.04% |

**Supplementary Table 2** Hydrogen bond occupancy (%) formed by the wildtype residue, E498, and mutated residue, A498 within neighboring residue during 500 ns of simulations.

| E498 | | |
| --- | --- | --- |
| Donor | Acceptor | H-bond occupancy (%) |
| S564-Main | E498-Side | 23.51% |
| E498-Main | S564-Main | 32.36% |
| S563-Side | E498-Side | 0.48% |
| S488-Side | E498-Side | 21.75% |
| S563-Side | E498-Main | 1.84% |
| H597-Side | E498-Main | 0.04% |
| A498 | | |
| A498-Main | S564-Main | 25.11% |
| S563-Side | A498-Main | 0.64% |

**Supplementary Table 3** Hydrogen bond occupancy (%) formed by the wildtype residue, R499, and mutated residue, G499 within neighboring residue during 500 ns of simulations.

| R499 | | |
| --- | --- | --- |
| Donor | Acceptor | H-bond occupancy (%) |
| R499-Main | R510-Main | 44.57% |
| R499-Main | T513-Main | 46.94% |
| R499-Side | S589-Side | 1.24% |
| R510-Main | R499-Main | 9.49% |
| R499-Side | E629-Main | 1.44% |
| R499-Side | H565-Side | 2.56% |
| T470-Side | R499-Main | 15.46% |
| R499-Side | E501-Side | 23.27% |
| R499-Side | D630-Side | 33.44% |
| R499-Side | D456-Main | 29.28% |
| R499-Side | N543-Side | 29.56% |
| R499-Side | F588-Main | 0.32% |
| R499-Side | D456-Side | 3.68% |
| R499-Side | E567-Side | 0.28% |
| R499-Side | H512-Side | 1.28% |
| R499-Side | E629-Side | 0.12% |
| R499-Side | M500-Main | 3.04% |
| G499 | | |
| G499-Main | R510-Main | 29.44% |
| R510-Main | G499-Main | 32.12% |

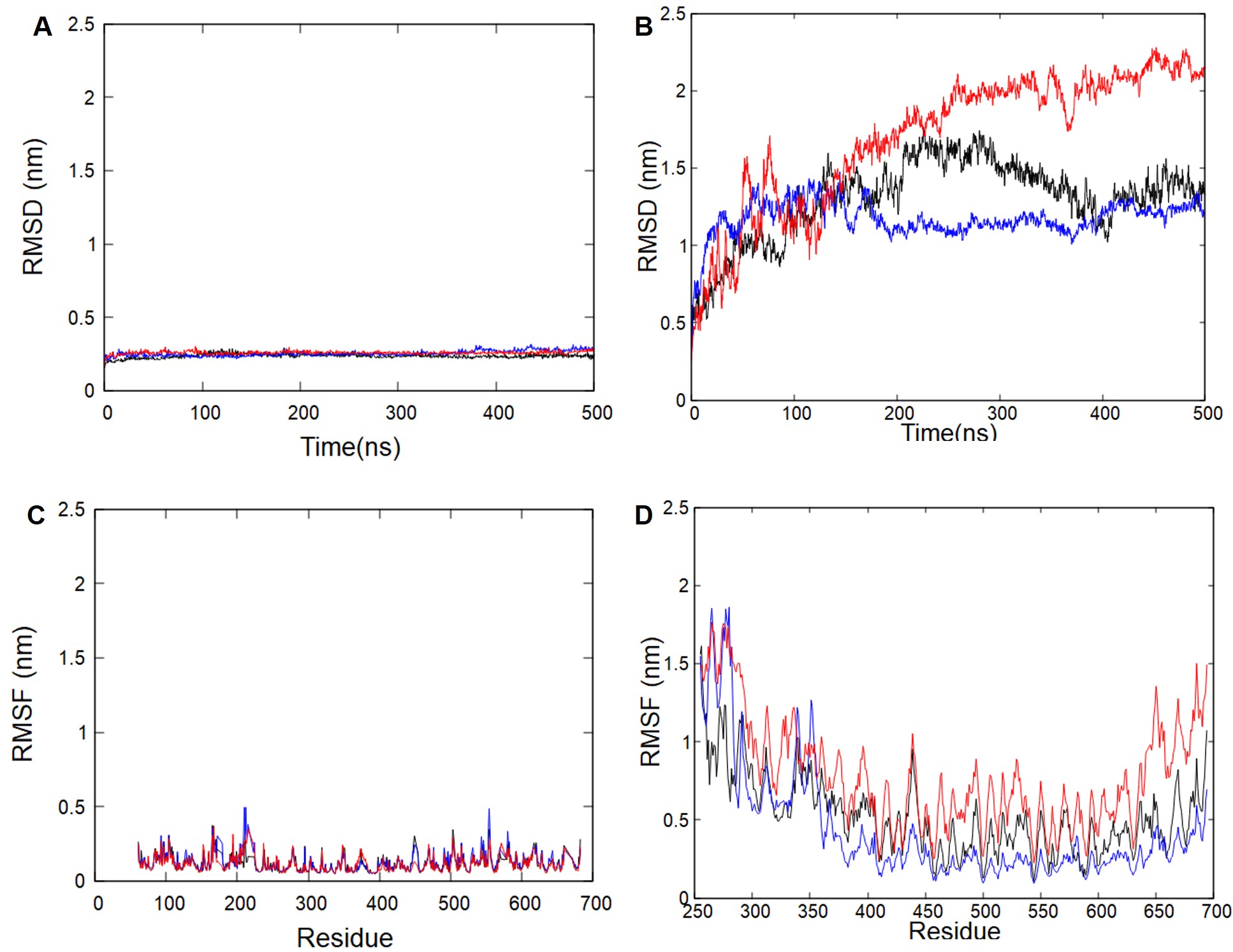

**Supplementary Figure 2 Replication analysis of LDLR-PCSK9 complexes.** RMSD of PCSK9 (A) and LDLR (B) and RMSF of PCSK9 (C) and LDLR (D) within 500 ns of simulations. Black (WT complex), blue (E498A complex), and red (R499G complex).
